# Bringing astrocytes into the spotlight of electrical brain stimulation

**DOI:** 10.1101/2024.02.28.582466

**Authors:** Miguel Aroso, Domingos Castro, Sara Costa Silva, Paulo Aguiar

## Abstract

Astrocytes, primarily viewed only as supportive units, are now emerging as active players in the information processing of the brain. Accumulated evidence supports that the bidirectional communication between astrocytes and neurons maintains complex animal behaviours such as memory formation and decision-making. The lack of characterisation of astrocytic electrophysiology is, in our opinion, associated with the early idea of a passive electrical nature of astrocytes, in opposition to the electrically active neurons. A better understanding of the effect of electrical stimulation on astrocytes’ physiology and activity will greatly strengthen the current knowledge in neural biology. Here, we assessed if astrocytes may have a role in therapies based on electrical brain stimulation by being able to respond to the same electrical stimulus used to modulate neuronal activity. To do so, we took advantage of microelectrode arrays (MEAs) capability to simultaneously record and deliver extracellular electrical signals. Additionally, we synchronized the recording of electrophysiological data with the recording of calcium activity, a hallmark of astrocytic activity. Here, we show that astrocytes respond to electrical stimulation with the generation of strong membrane voltage oscillations and simultaneous production of calcium waves, demonstrating, unequivocally, that astrocytes respond to electrical stimulation in the same range as neurons do. Importantly, these responses are dependent on the stimuli amplitude. Furthermore, membrane voltage oscillations are significantly reduced in the absence of extracellular calcium, but not abolished, while calcium activity is not detected.

## Introduction

The development of electrical brain stimulation therapies (namely deep brain stimulation, DBS) opened a new window for the treatment of neurological diseases. However, the mechanism of action of electrical brain stimulation is still far from being understood [2,3]. Neurons have been in the spotlight so far, but new findings point to the involvement of astrocytes as well [4,5]. Astrocytes, primarily viewed only as supportive units, are now emerging as active players in the information processing of the brain [6–8]. Astrocytic activity can be observed via changes in intracellular calcium dynamics. Those calcium events have characteristic spatial and temporal properties, which can last for several seconds, and be confined either to a single cell or propagate intercellularly, generating calcium waves that travel across many astrocytes [9]. Thus, astrocytic activity is generally assessed via calcium imaging. On the other hand, the electrophysiological activity of astrocytes is still a matter of debate [10]. Astrocytes do not fire action potentials, but they are not electrically silent cells. Indeed, several authors have described a plethora of different membrane signals on astrocytes [9]. Many of those membrane depolarisations were measured on astrocytes by patch-clamp (potassium, glutamate and GABA currents) in response to neuronal activity [11], which is accompanied by calcium signals. In addition, primary cultures were used to measure spontaneous electrophysiological activity in astrocytes [12,13] and to record high-frequency voltage oscillations as a result of electrical stimulation of astrocytic cells [14]. At this point, a clear relation between the electrophysiological and calcium activity in astrocytes is still missing.

Here, we assessed if astrocytes are equipped to play a role in DBS by being able to respond to the same electrical stimulus used to modulate neuronal activity. To do so, we took advantage of microelectrode arrays (MEAs) capability to simultaneously record and deliver extracellular electrical signals. Additionally, we synchronized the recording of electrophysiological data with calcium imaging, the hallmark of astrocytic activity. To clearly disconnect astrocytic response to electrical stimulation from a secondary type of response via stimulation of neurons we decided to work with rat primary astrocytic cultures. Although astrocytes are thought to be electrically silent, we reveal that they respond to electrical stimulation with the generation of membrane voltage oscillations (small membrane potential deflections) and the simultaneous production of calcium waves, demonstrating that astrocytes respond to electrical stimulation in the same range as neurons do. A better understanding of the effect of electrical stimulation on astrocytes’ physiology and activity will greatly strengthen the current knowledge in neural biology and enhance the translation of research findings into better devices and protocols for DBS.

## Results

### Electrical stimulation of astrocytes evokes/triggers calcium and membrane voltage oscillations

In the past years, some evidence has arisen pointing to the excitability of astrocytes via direct electrical stimulation, however, so far only the electrophysiological response has been investigated [14–16]. It is yet to be demonstrated how electrical stimulation is related to calcium intracellular fluctuations, the hallmark of astrocytic activity. We combined a microelectrode array (MEA) system and live-cell ready microscopy equipment to co-register both electrophysiological and calcium activity of astrocytes in response to direct electrical stimulation. We have used the ThinMEA chip (Multichannel Systems) for its 180μm thin recording area (similar to a coverslip) that facilitates the acquisition of microscopy imaging data. We started by testing a typical electrical pulse used to elicit neuronal activation, a biphasic square pulse of 200 μs duration and ±800 mV amplitude. As is observed in Figure 1, there is a generation of a calcium wave and membrane potential oscillations after the application of the stimuli. In this way, we can demonstrate undoubtedly that astrocytes respond to direct electrical stimulation.

**Figure 1.**
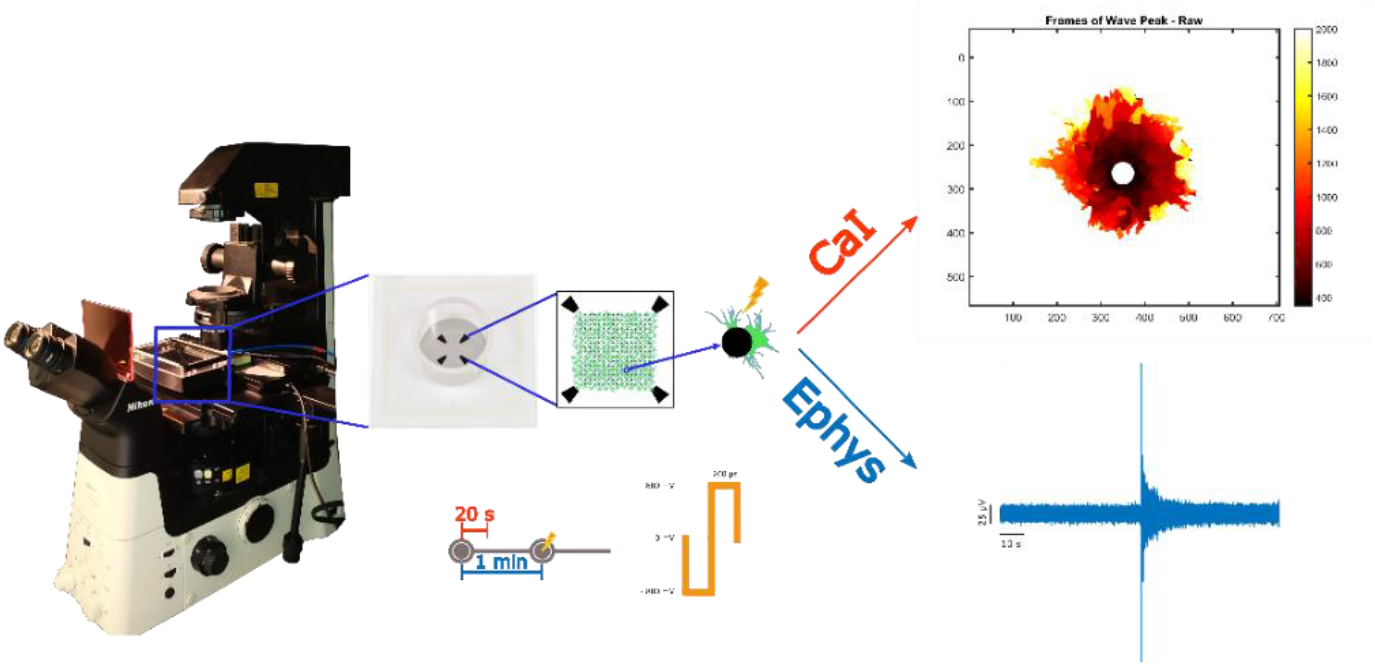
Illustration of the protocol used for the simultaneous acquisition of calcium imaging and electrophysiological data after electrical stimulation.

### Astrocytes respond to electrical stimulation in the same range as neurons do

Neuronal cultures, prepared on top of planar microelectrode arrays, are typically stimulated electrically within the range of 400 to 1000 mV [17]. To determine if astrocytes respond to this same range we tested the following voltage amplitudes: 400, 500, 600 and 800mv. The order of the stimuli was shuffled between rounds of stimulation. Since the measured calcium and electrophysiological signals have different natures we have to look at them differently. In this way, we have calculated the Fast-Fourier Transform (FFT) of the electrophysiological signal after stimulation and identified a strong increase in the power of the signal up to 100 Hz. This increase is dependent on the magnitude of the stimulus (Figure 2A). We then quantified the electrophysiological response as the fold change between the root mean square (RMS) of the signal and the average value of the baseline. The RMS of the response signal corresponds to the average RMS of the recorded voltage after each stimulus, while the baseline corresponds to the signal recorded before the start of the stimulation protocol. In the case of the calcium imaging data, since there is no baseline signal to compare with, we evaluated the probability of calcium wave initiation after each stimulus. This was achieved by dividing the number of calcium wave initiations by the total number of applied stimuli (Figure 2B).

**Figure 2.**
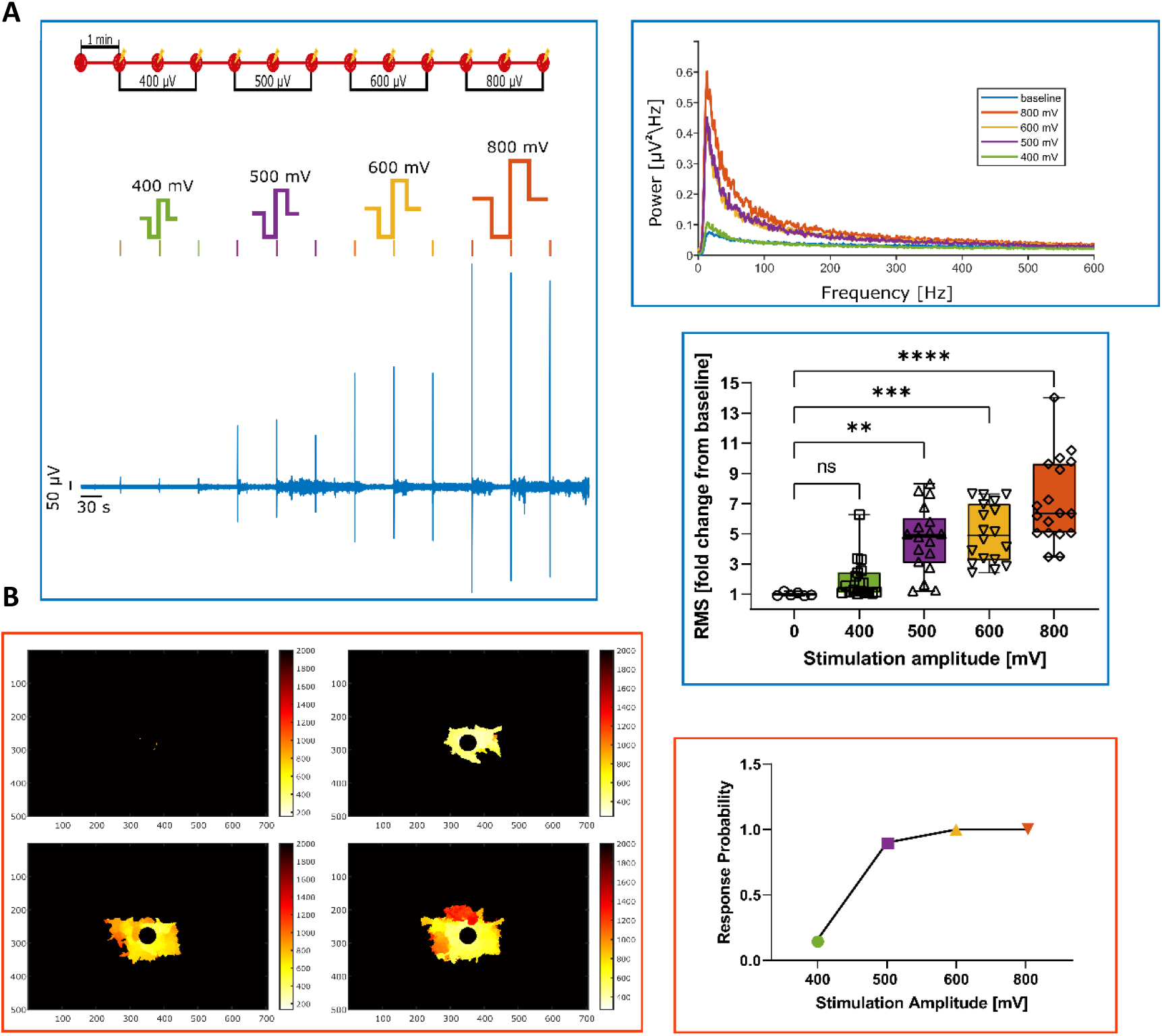
Evaluation of the response of astrocytes to increased voltage stimuli amplitude, ranging from ±400 mV up to ±800 mV. **A**. Analysis of the electrophysiological response to electrical stimulation by calculating the Fast Fourier Transform and the root mean square. **B**. Analysis of the calcium activity response probability to electrical stimulation. ns P>0.05, ^*^ P<0.05, ^**^ P<0.01, ^***^ P<0.001, ^****^ P<0.0001.

Stimuli of 400 mV generated very few times a calcium or a significant electrophysiological response of astrocytes to electrical stimulation. Above 500 mV the response became quite consistent in both cases. Thus, we determined 500 mV as the minimum amplitude necessary to consistently obtain a response of astrocytes to electrical stimulation. This is consistent with what is observed for neuronal cultures under the same conditions, meaning that the same electrical stimuli can excite both cellular populations.

### Electrically evoked calcium signals are faster, larger and stronger than spontaneous signals observed *in vitro*

Astrocytic cultures show spontaneous calcium signals that span from localised signals in microdomains to wider calcium waves. We developed an algorithm in MATLAB to extract the peak velocity of the travelling calcium signal, the area of propagation, the maximum intensity and the tau decay of the fluorescence signal for both spontaneous and stimulated calcium signals (800mV). It is important to note that the spontaneous signal was only measured during the baseline video, which was captured before the start of the stimulation protocol. As observed in Figure 3A, the peak velocity of the stimulated signal is 1.5 times faster than the spontaneous one, the maximum amplitude is over 2 times higher, the propagation area is one order of magnitude higher and the tau decay is almost 1.5 times higher. It is evident from the results that the calcium signal obtained through electrical stimulation is much stronger than the signals observed spontaneously. This indicates that astrocytes not only respond to electrical stimulation but also that it might have a profound impact on their activity. This is further supported by the fact that the astrocytic response is weakened with the consecutive stimulation of cells. In Figure 3B is possible to observe that all parameters of the calcium wave evaluated, except for tau decay, decrease in the second and third stimuli. We chose to stimulate astrocytes with 1-minute intervals because, in the preliminary tests, this seemed to be enough time for the signal to get back to baseline, both when looking at the calcium and the electrophysiological signals. The large response of astrocytes may prevent them from fully restoring calcium levels for the next event, leading to a decline in response to electrical stimulation. A similar refractory period was observed in vivo, where astrocytes had a significantly diminished response to a second event [18] and ex-vivo where a decrease in average calcium amplitude was observed after stimulation of astrocytes with baclofen [19].

**Figure 3.**
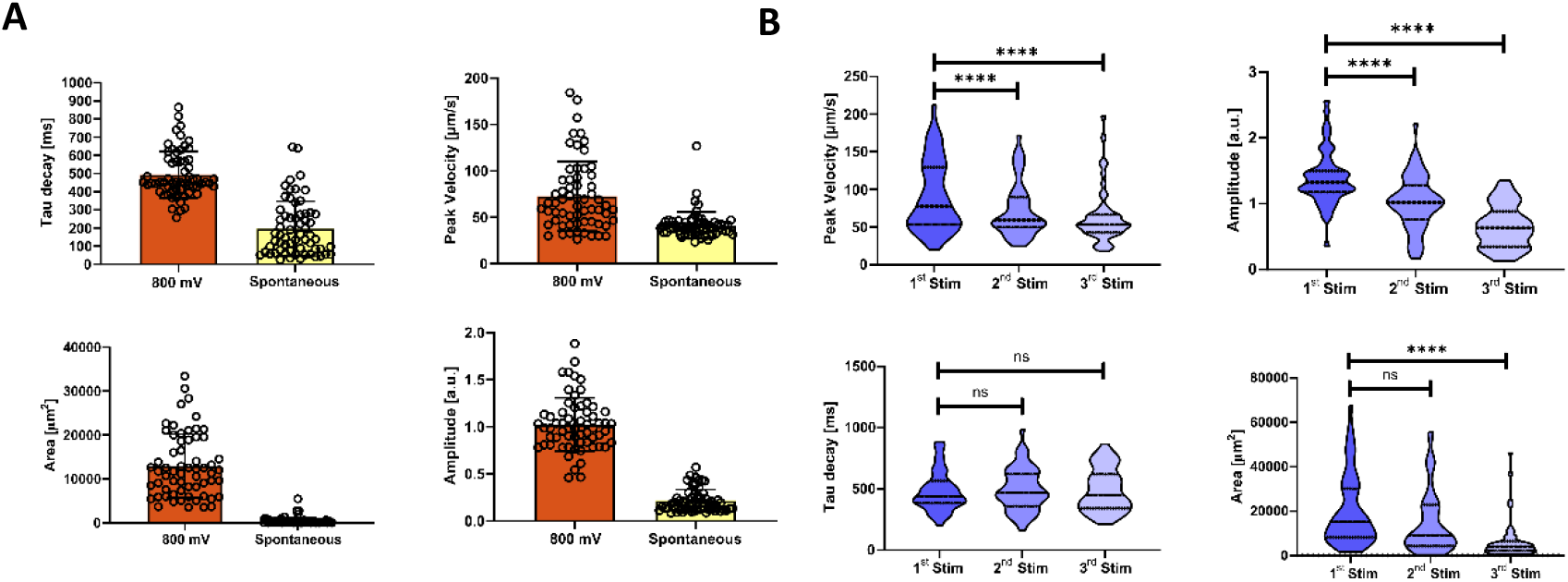
Characterization of the calcium wave dynamics. **A**. Comparison between evoked (±800 mV) and spontaneous calcium signals. **B**. Evaluation of the calcium wave parameters that emerge from the response of astrocytes to 3 consecutive stimuli of ±800 mV separated by 1 min. ns P>0.05, ^****^ P<0.0001

### The astrocytic response exhibits/displays electrical signals even in the absence of a calcium signal

To determine the dependence of astrocytic activity on extracellular calcium, we first stimulated the cells to confirm the excitability of the culture and then substituted the extracellular solution with a calcium-free solution. After the removal of calcium, we did not detect calcium activity on astrocytes upon stimulation, however, it was still possible to record electrophysiological signals (Figure 4). There was a reduction in the magnitude of the electrophysiological signal to around 50% of the control. This indicates that astrocytic calcium activity in response to electrical stimulation is only partially dependent on the presence of extracellular calcium. This partial dependence suggests a mechanism that relies on Ca^2+^-sensitive ryanodine receptors (RYR): extracellular calcium triggers RyR activity by increasing intracellular calcium concentration that leads to Ca^2+^ release from the endoplasmic reticulum (ER) [20,21]. Furthermore, it also indicates that the entry of calcium in astrocytes via ionic channels contributes to the membrane voltage oscillations but they are not the sole contributors. On the other hand, Ca^2+^ channels become permeable to Na^+^ and K^+^ in the absence of extracellular Ca^2+^ [21], which might contribute to the recorded electrophysiological activity.

**Figure 4.**
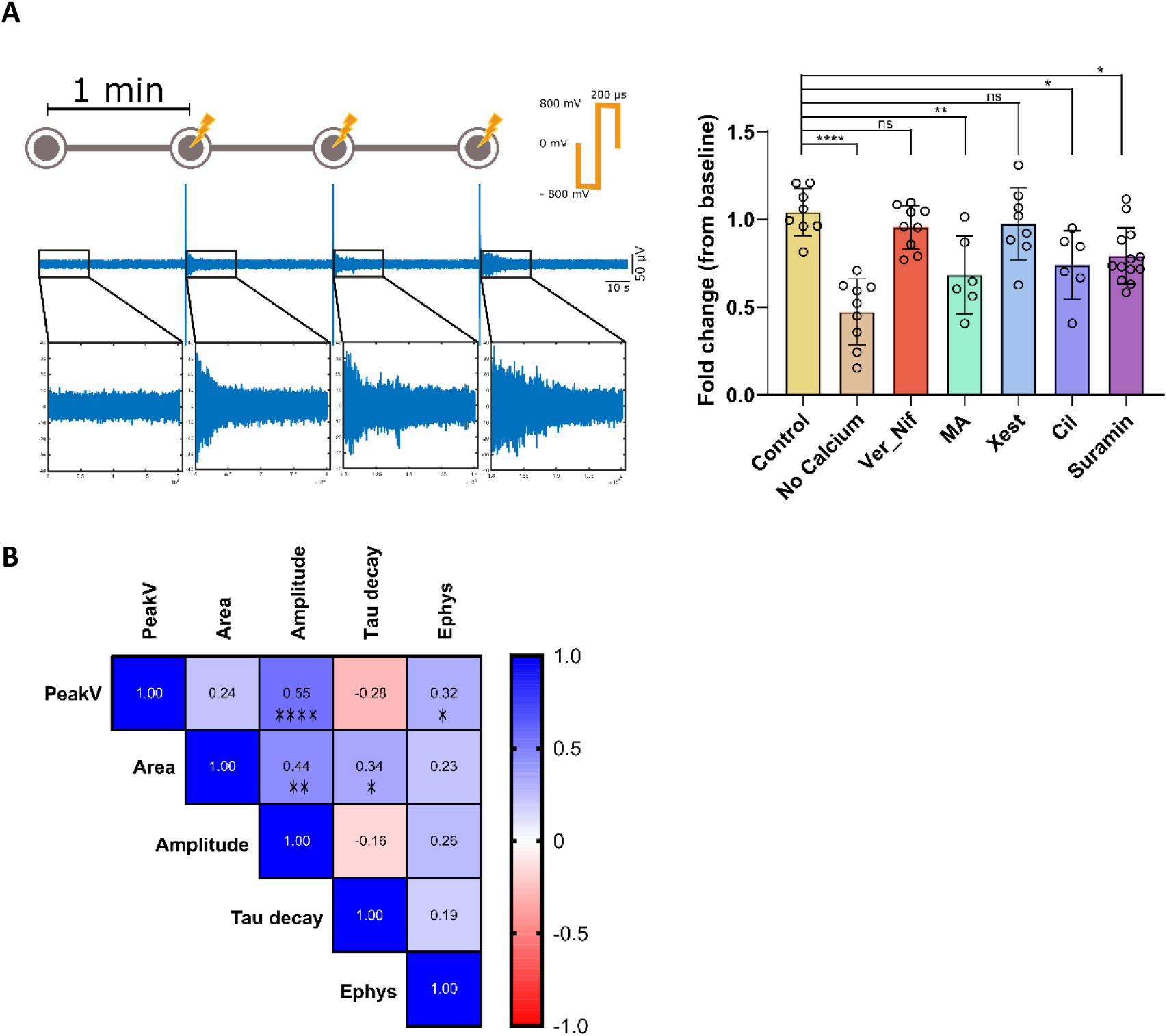
Mechanisms involved in the electrophysiological response of astrocytes to electrical stimulation. **A**. Evaluation of the of pharmacological treatment in the electrophysiological response of astrocytes to 3 consecutive stimuli of ±800 mV separated by 1 min. **B**. Pearson correlation between the RMS of the electrophysiological response and calcium wave parameters of astrocytes to a stimulus of ±800 mV. ns P>0.05, ^*^ P<0.05, ^**^ P<0.01, ^***^ P<0.001, ^****^ P<0.0001.

### Electrically evoked electrophysiological signals are decreased by inhibition of gap junctions, P2X receptors and N-type Voltage-Gated Calcium Channels

In the course of this work, we realized that the calcium signalling in astrocytes in response to electrical stimulation is not consistent across multiple stimuli. On average, the second stimulus gives rise to a smaller calcium wave and a third stimulus further reduces this response. To analyse the effect of a pharmacological treatment on the dynamics of the calcium wave we have to measure the response of astrocytes to electrical stimulation before and after the treatment. But if the response changes in consecutive stimuli it is not possible to distinguish if a change in the response after pharmacological treatment is due to the addition of the drug or due to different responses from the cells to the same stimuli. So, we have focused on the electrophysiological data to evaluate the astrocytic response to electrical stimulation after pharmacological treatment.

We started by investigating if voltage-gated calcium channels (VGCC) are involved in the electrophysiological response of astrocytes to electrical stimulation we blocked VGCC with Verapamil (L-type) and Cilnidipine (L and N-type). We did not observe a significant alteration in the magnitude of membrane voltage fluctuations in the case of inhibition with Verapamil. On the other hand, inhibition with Cilnidipine resulted in a 30% reduction in the signal magnitude. This indicates a significant role of N-type VGCC in the transduction of the electrical stimuli and entry of calcium into the cells, however, this is not the only mechanism associated with the origin of membrane voltage oscillations.

There are several mechanisms involved in the propagation of calcium waves, namely IP3 diffusion via gap junctions or extracellular diffusion of ATP [22]. To investigate if these mechanisms are involved in the propagation of calcium waves after electrical stimulation of astrocytes we have used the following inhibitors: Meclofenamic acid (MA) to inhibit gap junctions; Xestopongin C (XestC) to inhibit IP3 receptors and Suramin to inhibit ATP receptors (P2X) at astrocytic membranes. Both MA and Suramin had a significant impact on the measured RMS signal with a 33% and 30% reduction, respectively. Whereas XestC did not affect significantly the magnitude of the astrocytic electrophysiologic signals. We have to keep in mind that the electrode is only able to receive electrical signals from the close surroundings of the electrode (≈50 μm radius) and the calcium wave travels up to 200 μm away from the start point, which corresponds to the position of the electrode since this is the point of stimulation. This means that we are analysing the effect of the inhibitors on the calcium wave propagation only on the astrocytes close to the electrode. On the other hand, astrocytes form a syncytium of interconnected cells that is thought to equilibrate the membrane potential across those cells providing a strong electrical coupling [23,24]. Our results indicate that at least blocking gap junctions or ATP membrane receptors reduce locally the magnitude of membrane voltage oscillations, indicating that both mechanisms might be relevant for the propagation of calcium waves.

To explore if there is a linear relationship between the measured calcium signal and membrane voltage oscillations we calculated the Pearson correlation coefficient between the RMS obtained for the electrical signal and each of the metrics (Peak velocity, amplitude, propagation area and tau decay) obtained for the calcium data. Only peak velocity and RMS were found to be moderately positively correlated (r = 0.32, p < 0.5).

## Discussion

In the past decades, electrical brain stimulation therapy has been successfully used to treat neurological diseases, like Parkinson’s disease [25] and epilepsy, and even psychiatric disorders, like obsessive compulsion disorder or depression [26]. However, these therapeutic approaches are constrained by side effects whose source is not understood [27,28] and their efficacy cannot be uniformly applied to the full spectrum of patients [28]. In the last years, major improvements have been made in the surgical technique [29] while, in comparison, there has been virtually no advance in the stimulation protocols since its implementation [28,30]. To increase the efficacy of the treatments (e.g. DBS) it is necessary to dissect the effect of electrical stimulation on the cellular populations of the brain. Here, we set to unambiguously reveal the astrocytic response to electrical stimulation in the same range used to excite neurons.

The excitability of astrocytes via intracellular calcium fluctuations has been demonstrated from in vitro up to in vivo settings across the years [31]. There are various mechanisms involved in the origin of the calcium signal, such as Gq-coupled receptor-induced Ca^2+^ signalling, reversed operation of the Na^+^/Ca^2+^ exchanger (NCX), P2X purinoreceptors, ionotropic glutamate receptors (AMPA and NMDA), transient receptor potential (TRP) cation channels. [9,32] or voltage-gated calcium channels (VGCC). On the other hand, astrocytic membranes’ voltage oscillation is less recognized, but they arise spontaneously [12,13] and in response to electrical stimulation [14] Whether those voltage oscillations are related to calcium signalling (astrocytic activity) has not yet been demonstrated.

We used a specific type of MEA chip (ThinMEA) to simultaneously record calcium imaging and electrophysiological data. This allowed us to clearly show that electrical stimulation of astrocytes evokes/triggers calcium and membrane voltage oscillations, demonstrating that those appear in response to electrical stimulation and that astrocytes are electrically excitable. The evoked calcium response propagates radially to the point of stimulation and can reach as far as up to 200 μM. This is in line with what was described before [33–35]. The observed membrane voltage oscillations are characterized by fast and long-lasting electrical signals with a frequency of up to 100 Hz. However, Fleischer and colleagues described extracellular voltage oscillations that cover a frequency spectrum up to 600 Hz [14]. One main contributor to these differences is the fact that they have applied a high-pass filter with a cut-off frequency of 100 Hz in the acquisition hardware, thus refraining from detecting any signal below 100 Hz.

Besides demonstrating astrocyte responsiveness to electrical stimulation, it is also important to confirm whether they respond to the same range of stimulation used to excite neurons. After analysing calcium imaging and electrophysiological data, we have consistently observed astrocytes to be excitable above ±500 mV. This is very similar to what is observed in neuronal cultures under the same conditions [17]. In this way, we believe that the stimulation protocol used to modulate neuronal activity might as well change the activity dynamics of the astrocytic population. This is further supported by the fact that the calcium activity evoked by the electrical stimulation is much stronger than the spontaneous activity observed in these cultures. The average peak propagation speed of the calcium wave, measured after electrical stimulation was close to 60 μm/s. This is in agreement with previous results describing, in vivo, the mean speed propagation of 61 ± 22 μm/s [36]. Of note, it is not possible to do the same comparison for the electrophysiological data because we have not been able so far to detect spontaneous astrocytic electrical activity with our MEA system.

We used pharmacological manipulation to explore the basic mechanisms behind the generation of the astrocytic response to electrical stimulation but came across a significant constraint regarding the calcium imaging data. We set a protocol that relies on 3 stimuli, each separated by 1-minute intervals to allow for cell recovery. Indeed, we observed that after 1-minute the generated calcium wave was over and the cellular membrane potential had already returned to baseline. However, in the case of the calcium wave, the second and third stimuli most often generated a significantly smaller response in terms of peak velocity, fluorescence intensity and total area propagation (Figure 3B). This might be associated with a refractory period of calcium signalling in astrocytes that may become evident after the massive calcium release observed after stimulation. Fedotova and colleagues recently proposed an astrocytic refractory period that arises due to the depletion of intracellular calcium stores in astrocytes, which affects consecutive calcium signals [18]. This excludes the use of calcium imaging data to clarify the mechanisms behind its dynamics as the response is inherently different when comparing the first stimuli (baseline) with the response obtained after pharmacological manipulation. Nevertheless, it was still possible to evaluate the effect of the removal of extracellular calcium from the recording solution on the astrocytic response to stimulation, since it resulted in the complete abolishment of any measurable calcium signal on astrocytes after electrical stimulation. Regarding the electrophysiological data after the removal of extracellular calcium, there was a marked reduction in the recorded signal. This indicates that calcium signalling in astrocytes, in response to electrical stimulation, is dependent on extracellular calcium but not the associated membrane voltage oscillations. The movement of calcium ions across the membrane may have a significant contribution to changes in membrane voltage, but other ions should also be involved in the process of generation of membrane voltage oscillation, for example, calcium channels become less selective in the absence of extracellular calcium [21].

Contrary to calcium imaging, the measured electrophysiological astrocytic response to stimulation is very stable across stimuli, thus we focused on this data to evaluate the response of astrocytes under pharmacological manipulation. It should be noted that it is only possible to measure the response of astrocytes located in close proximity to the stimulating electrode, which means that changes in the membrane potential of cells that are further away from the electrode and participate in the propagation of the calcium wave might not be directly detected. However, astrocytes form a syncytium of interconnected cells that provides a strong electrical coupling that balances the membrane potential across several cells [23], opening the door to long-distance measurement of membrane potential oscillations. We tested chemical antagonists for gap junctions (Meclofenamic acid), P2X receptors (Suramin), VGCC (Verapamil and Cilnidipine) and IP3 receptors (IP3R) (Xestopongin-C). A significant reduction in the magnitude of membrane voltage oscillations was observed after the addition of MA, Suramin and Cilnidipine. This indicates that gap junctions and P2X receptors are involved in the spread of membrane voltage oscillations and, most probably, in calcium wave propagation. The inhibition of VGCC revealed a reduction in the membrane potential oscillations with Cilnidipine, pointing to the involvement of N-type channels. Previously a dose response to Cilnidipine was observed by Fleischer and colleagues that led to a decrease in high-frequency oscillation (HFO) power in response to electrical stimulation [14]. On the other hand, blocking IP3 receptors did not result in any measurable alteration in the electrophysiological signal. IP3 is generated intracellularly after Gq-coupled receptor activation and binds to IP3 receptors on the membrane of the endoplasmic reticulum promoting the release of calcium [37]. However, it is essential to consider that IP3R blockage may not have an electrophysiological readout. Therefore, ideally, calcium imaging should be performed to evaluate this, but as discussed before, it was not possible to quantify the effect of pharmacological manipulation via calcium imaging. Still, qualitatively it was possible to observe the generation of a calcium wave after blocking IP3R. This supports the involvement of other known mechanisms in the generation and propagation of a calcium wave, such as the stimulation of RYR after extracellular calcium entry in the cell [38].

At last, we assessed the potential correlation between the quantification derived from calcium imaging and electrophysiology. Only a moderate positive relationship (r = 0.32, p < 0.5) was observed between the measured peak velocity and the RMS of the electrophysiological signal. This means that measuring one signal is not enough to infer the features of another signal or vice-versa, pointing to the complementarity of both signals.

## Conclusion

There is mounting evidence suggesting that astrocytes may play a crucial role in the beneficial effects of electrical brain stimulation therapy, however, it was not clear so far if astrocytes respond directly to electrical stimulation in the same range as neurons do. We have shown here that astrocytes respond to the same electrical stimulus that evokes neuronal activity via intracellular calcium fluctuations and membrane voltage oscillations. Furthermore, the calcium response is characterized by the generation of a calcium wave, is dependent on the presence of extracellular calcium and is much larger than the detected spontaneous signals. Membrane voltage oscillations are significantly reduced in the absence of extracellular calcium, but not abolished, and are partially dependent on N-type VGCC, gap-junctions and ATP signalling. Additionally, a modest correlation was obtained between the peak velocity of the calcium wave and the RMS of the membrane voltage oscillations. This implies that the information provided by both signals is complementary rather than redundant. Future studies are expected to reveal the effect of astrocytic electrical stimulation on neuronal activity and help to understand the mechanisms behind deep brain stimulation, or even to uncover astrocytes as possible therapeutic targets for therapeutic electrical stimulation.

## Material and Methods

### Isolation and culture of astrocytes

All procedures conducted adhered to the institutional and European Guidelines for the ethical care and use of laboratory animals. Primary cultures of astrocytes were derived from the cortex of 1-to 2-day-old Wistar Han rat pups. Following decapitation, brains were promptly removed, and the cortex was meticulously dissected, excluding the meninges and cerebellum. The cortex tissue underwent mechanical dissociation and enzymatic digestion with 0.01% trypsin and 0.01% DNAse I (Sigma-Aldrich) at 37°C for 15 minutes. Subsequently, cells were seeded in 75cm2 flasks pre-coated with 10mg/mL poly-D-lysine (PDL) (Sigma-Aldrich) and maintained in culture using DMEM-Glutamine high glucose medium (Corning) with 10% Fetal Bovine Serum (FBS) (Capricorn) and 1% Penicillin-Streptomycin (Gibco) at 37°C and 5% CO2.

To obtain astrocytic cultures and mechanically detach microglia and oligodendrocyte precursor cells were mechanically detached. Flasks underwent shaking at 220rpm, 37°C for16h 12 days after initiation, followed by weekly shakes for two subsequent weeks. After the final shake, cells were trypsinized with 0.05% Trypsin-EDTA (Gibco) and plated at a 1:2 ratio. Astrocytes were used from the third passage (P) onward, until reaching P8.

### Calcium imaging

For Ca^2+^ imaging, astrocytes were trypsinized and seeded at a density of 2x105 cells on poly-D-lysine-coated (10μg/mL)(Sigma-Aldrich) ThinMEA (MultiChannel© systems) chips, where they were cultured for 2 days after seeding for cellular recovery. ThinMEAs have a thickness of only 180μm, making them the optimal choice for high-resolution imaging. These chips feature 252 microelectrodes, each with a diameter of 30μm and spaced at 200μm spacing, along with four reference microelectrodes, embedded in an ultra-thin glass substrate on a durable ceramic carrier. Previous to image acquisition and electrical stimulation cells were stained with the calcium indicator dye CAL-520-AM^®^ (AAT Bioquest). Cell medium was replaced by 1x Hepes-buffered Hanks balanced salt solution (HBSS) with 0,04% Kolliphor^®^ P407 (BASF) and 10μM CAL-520-AM^®^ dye, and incubated at 37°C, 5% CO2 for 30 min, followed by 10 minutes at room temperature. Before image acquisition the dye was removed and the solution substituted by fresh HBSS.

### Image acquisition

Astrocytic monocultures seeded on ThinMEAs underwent staining with the calcium dye CAL-520-AM immediately before start of the experiment. Images were acquired with a sCMOS camera Prime 95B, 22 mm (Teledyne Photometrics), mounted on a Nikon Eclipse Ti2-E inverted microscope (Nikon) with a Nikon Pl Apo 40X/1.15NA (water-immersion) objective. Time stacks were acquired at 100Hz with 2x2 binning for a total of 20 seconds (2000 frames).

### Electrophysiology recording and electrical stimulation

The cultures were kept at 37 °C and 5% CO2 using a stage-top incubator (ibidi GmbH) adapted to the headstage of the MEA2100-256 system (MultiChannel© systems). The electrophysiology signals were sampled at 10 kHz and a 0.1 Hz high-pass filter and a 3500 Hz low-pass filter were used in the MEA2100-256 before signal amplification. A single biphasic square stimulus ranging from ±400 mV up to ±800 mV, each phase lasting 200μs, was administered at the selected electrode.

## Acknowledgements

This work was supported by Prémio Mantero Belard, Santa Casa da Misericordia de Lisboa (grant MB-12-2022), and by Foundation ‘la Caixa’ -CaixaResearch Health 2022 (grant HR22-00189). Miguel Aroso was funded by FCT – Fundação para a Ciência e a Tecnologia, CEEC-IND contract CEECIND/03415/2017. Domingos Leite de Castro was funded by FCT, grant contract SFRH/BD/143956/2019. Sara Costa Silva was funded by FCT, grant contract 2020.05463.BD.

## Author contributions

M.A. contributed to the methodology, experiments, formal analysis, visualization, and writing of the original draft. S.C.S. contributed to experiments and formal analysis. D.C. contributed to formal analysis. P.A. contributed to conceptualization, methodology, funding, and work supervision. All authors contributed to the review and editing of the final manuscript.

## Competing Interests

The authors declare no competing interests.

## Data Availability

The data that support the findings of this study are available from the corresponding author upon reasonable request.

## Notes

### Competing Interest Statement

The authors have declared no competing interest.

